# Deubiquitinase JOSD1 tempers hepatic proteotoxicity

**DOI:** 10.1101/2024.07.03.601825

**Authors:** Saheli Chowdhury, Abhishek Sen, Debajyoti Das, Partha Chakrabarti

## Abstract

Derangements in protein homeostasis and associated proteotoxicity mark acute, chronic, and drug-induced hepatocellular injury. Metabolic dysfunction-associated proteasomal inhibition and the use of proteasome inhibitors often underlie such pathological hepatic proteotoxicity. In this study, we sought to identify a candidate deubiquitinating enzyme (DUB) responsible for reversing the proteotoxic damage. To this end, we performed a siRNA screening wherein 96 DUBs were individually knocked down in HepG2 cells under proteasomal inhibitor-induced stress for dual readouts, apoptosis, and cell viability. Among the putative hits, we chose JOSD1, a member of the Machado-Josephin family of DUBs that reciprocally increased cell viability and decreased cell death under proteotoxicity. JOSD1-mediated mitigation of proteotoxicity was further validated in primary mouse hepatocytes by gain and loss of function studies. Marked plasma membrane accumulation of monoubiquitinated JOSD1 in proteotoxic conditions is a prerequisite for its protective role while the enzymatically inactive JOSD1 C36A mutant was conversely polyubiquitinated, does not have membrane localization and fails to reverse proteotoxicity. Mechanistically, JOSD1 physically interacts with the suppressor of cytokine signaling 1 (SOCS1), deubiquitinates it and enhanced its stability under proteotoxic stress. Indeed, SOCS1 expression is necessary and sufficient for the hepatoprotective function of JOSD1. *In vivo*, adenovirus-mediated ectopic expression or depletion of JOSD1 in mice liver respectively protect or aggravate hepatic injury when challenged with proteasome blocker Bortezomib. Our study thus unveils JOSD1 as a potential candidate for ameliorating hepatocellular damage in liver diseases.

## INTRODUCTION

Liver is the major protein metabolic hub and dysregulated hepatic protein turnover could adversely incriminate to a range of liver diseases. Accumulation of ubiquitinated proteins and appearance of classic Mallory Denk Bodies in liver histology across many chronic liver diseases like ASH, NASH, Wilson’s disease, HCC, etc. support towards the conspicuous involvement of altered protein homeostasis in these conditions^1^. Moreover, disruptions in protein homeostasis cause ER stress and give rise to hepatic insulin resistance associated with diminished proteasome activity^2^. There is a remarkable accumulation of ubiquitin with a complimentary drop in the proteasome activity in livers of high fat diet fed mice. A subset of human NASH patients also show genetic signature consistent with reduced proteasome activity^3^. Few proteasomal inhibitors including Bortezomib are used widely chemotherapy in multiple myeloma patients with observable but tolerable hepatotoxicity^4,5^. Clinical cases are quite prevalent wherein multiple myeloma patients treated with Bortezomib display significant hepatocellular damage and injury ^6,7,8^.

The ubiquitin proteasome system comprising of Ubiquitin, E1-Ubiquitin Activating Enzyme, E2-Ubiquiting Conjugating Enzyme, E3 Ubiquitin Ligase, 26s proteasome and Deubiquitinating Enzymes (DUBs) constitute major cellular proteolytic machinery. By removing ubiquitin from the cognate substrates DUBs curtail undue or excessive protein degradation^9^, turn off or turn on a signalling pathway by proofreading cellular ubiquitin adducts, protect activated ubiquitin from nucleophilic attack^10^, liberate unanchored ubiquitin chains from 26s proteasome and recycle the cellular ubiquitin pool. Humans harbour roughly around 100 DUBs^11,12^ and they are divided into five different families, four of which are cysteine proteases and the fifth family constitutes zinc-dependent metalloproteases. Ubiquitin specific processing proteases (USP, or UBP in yeast), the ubiquitin C-terminal hydrolases (UCH), ovarian tumor related proteases (OTU), and the Josephin/Machado-Joseph disease proteases (MJD) constitute the cysteine protease DUB family while JAB1/MPN/Mov34 metalloenzyme (JAMM) comprise the zinc-dependent metalloprotease DUB family^11^. USP, UCH, OUT and MJD families have a conserved cysteine and histidine boxes with Cys, His and Asp as the residues in the active site catalytic triad whereas the JAMM family contains two conserved His and Asp residues coordinated to a Zn residue in its active site^13^. Certain DUBs are specific for selected types of ubiquitination while others exhibit a broad spectrum of action. The role of DUBs in chronic liver diseases is recently being unravelled. Cylindromatosis (CYLD) and TNFAIP3 are DUBs which have been implicated in hepatic apoptosis, inflammation and fibrosis in the context of non-alcoholic fatty liver disease and hepatocellular carcinoma^14,15,16^. Also, USP18 when overexpressed specifically in HFD fed mice livers could improve hepatic steatosis and insulin sensitivity through deubiquitinating TAK1^17^. USP2 is known to regulate hepatic gluconeogenesis and glucose metabolism by modulating hepatic expression of 11β-hydroxysteroid dehydrogenase 1^18^.

The cysteine protease JOSD1 belongs to the MJD family of DUBs that primarily cleaves K48 ubiquitin chains when it is monoubiquitinated and has its catalytic active site at C36^19,20^. JOSD1 is known to localise to the plasma membrane and interact with actin cytoskeleton to regulate membrane dynamics, cell motility and influence endocytosis^19^. JOSD1 can inhibit IFN-1 induced signalling pathway and antiviral response by physically interacting with SOCS1 and enhancing SOCS1 stability by cleaving K48-ubiquitin chains from SOCS1^21^. JOSD1 could also deubiquitinate MCL1 and stabilize MCL1 to suppress mitochondrial apoptosis and confer chemoresistance in gynaecological cancer^20^. The expression of JOSD1 was aberrant in HNSCC specimens and its depletion improved cisplastin-induced apoptosis of HNSCC cells, suppressed tumour growth and improved chemosensitivity *in vitro* via regulation by the epigenetic regulator, BRD4^22^. JOSD1 deubiquitinates and stabilises mutant JAK2 (JAK2-V617F) and therefore inactivation of JOSD1 via small molecules led to degradation of mutant JAK2 in MPNs providing a novel therapeutic approach for the disease^23^. JOSD1 was overexpressed in lung adenocarcinoma (LUAD) tissues and its knockdown suppressed tumour cell proliferation and inhibited metastasis. It was seen that JOSD1 could deubiquitinate and stabilise Snail protein which promoted epithelial to mesenchymal transition (EMT) in LUAD^24^. In Duck Tembesu virus infection, JOSD1 interacts with SOCS1 via its SH2 domain and stabilises SOCS1. SOCS1, in turn, acts as an E3 ubiquitin ligase mediating the degradation of IRF7 and inhibiting type I interferon production leading to the proliferation of the virus^25^. MCL-1 stabilization by JOSD1 has also been implicated in conferring radioresistance in oral squamous cell carcinoma (OSCC) wherein TRAF4 mediates MCL-1 phosphorylation and making it inaccessible for JOSDI interaction^26^. However, the role of JOSD1 in liver diseases and particularly in proteotoxicity is unknown.

In this study, we identified JOSD1 as a hitherto unknown candidate for mitigating and combating hepatic proteotoxicity. Using various *in vitro* and *in vivo* models of hepatic proteotoxicity, we showed that JOSD1-mediated deubiquitination and stabilization of SOCS1 is the putative molecular basis of hepatoprotection. Thus, we discovered a novel role of JOSD1 in ameliorating hepatic apoptosis under conditions of proteotoxicity.

## MATERIALS AND METHODS

### Animal Study

Protocols for animal experiments were approved by Institutional Animal Ethics Committee at CSIR-IICB under the aegis of Committee for Control and Supervision of Experiments on Animals (CPCSEA), Ministry of Environment and Forest, Government of India. 6-8 weeks old C57BL/6 male mice were divided into three groups. One group was injected with 3X10^11^ pfu AdEGFP and the other two with 3X10^11^ pfu shJOSD1 and 3X10^11^ pfu AdJOSD1 respectively via tail vein. After a week, each group was intraperitoneally injected with 5 mg/kg of Bortezomib (Merck, Burlington, Massachusetts, United States of America) and sacrificed after 16 h. Serum was collected for liver enzyme analysis and liver sections were collected in formalin (for histological analysis) and lysis buffer (for protein lysate preparation for Western Blot).

### AST estimation

Serum was collected from blood of sacrificed mice by cardiac puncture. Aspartate aminotransferase (AST) levels were measured using a commercially available kit (Randox Laboratories, Crumlin, County Antrim, United Kingdom) using the kinetic method following the manufacturer’s protocol. Values were obtained using a semi-auto analyser Microlab 300 (Clinical Systems, EliTech Group, Puteaux, France).

### Histology, Immunohistochemistry and Immunofluoroscence

Liver tissue sections were processed for H&E and images were taken using Evos XL Core (Thermo Fisher Scientific). Liver tissue sections were first deparafinized by heating at 85°C for 10 min followed by treatment with decreasing gradients of alcohol. The sections were then permeabilised and the antigens were retrieved by heating with Sodium Citrate Buffer. Immunohistochemistry was then performed using two kits: VECTASTAIN ABC KIT (Biotinylated Horseradish Peroxidase Anti-rabbit IgG) [VectorLabs, Newark, California, United States of America] and ImmPRESS Duet Double staining Polymer kit (Horseradish Peroxidase Anti-Mouse IgG/Alkaline Phosphatase Anti-Rabbit IgG) [VectorLabs, Newark, California, United States of America] following the manufacturer’s protocol. Images were taken with Evos XL Core (Thermo Fisher Scientific, Waltham, Massachusetts, United States of America). For Immunofluorescence, blocking buffer was added to block the non-specific binding of antibody. The sections were then incubated with Cleaved Caspase 3 primary antibody tagged with AlexaFluor 647 (Cell Signaling Technology, Danvers, Massachusetts, United States of America) at a dilution of 1:100 overnight in a humidified chamber at 4°C. The next day, the sections were stained with 25 µg/ml HOECKST342 (Thermo Fisher Scientific, Waltham, Massachusetts, United States of America) and taken for imaging Leica TCS SP8 STEDmicroscope (Leica Microsystems, Wetzlar, Germany).

### Generation of Adenovirus

shJOSD1 carrying adenovirus was generated using BLOCK-iT^TM^ Adenoviral RNAi Expression System (Thermo Fisher Scientific, Waltham, Massachusetts, United States of America) following the manufacturer’s protocol. The following sequences were used to generate the ds oligo of shJOSD1: Top strand oligo - 5’ caccgcacaagaagagcatgctgggaaatgggaacgaattcccatttcccagcatgctcttcttgtg 3’; Bottom strand oligo - 5’ aaaacacaagaagagcatgctgggaaatgggaattcgttcccatttcccagcatgctcttcttgtgc 3’. Briefly, the pENTR clone carrying the shJOSD1 ds oligo was obtained, recombined with pDEST, and the final product was digested with PacI enzyme and transfected in HEK293A cells to generate viable adenoviral particles. Crude adenoviral HEK293A cell lysate expressing JOSD1 was purchased from ABM (#252540540200) [New York, New York, United States of America]. Once the cells started to be lysed by actively replicating viral particles, the media along with the adherent cells were harvested and after repeated cycles of freezing and thawing, active viral particles were purified using Pure Virus^TM^ Adenovirus Purification Kit (Cell Biolabs, San Diego, California, United States of America) as per manufacturer’s protocol. The viral titre was determined spectrophototmetrically and the purified virus was aliquoted and stored at −80°C until further use.

### Primary mouse hepatocytes

Primary mouse hepatocytes were isolated and cultured as described previously^27^. Briefly, 8-12 weeks old chow-fed black male mouse (C57bl/6) was sacrificed, portal vein was cannulated, and liver was perfused with 20 ml of HBSS (Hank’s Balanced Salt Solution; 5mM KCl, 0.4mM KH_2_PO_4_, 4mM NaHCO_3_, 140mM NaCl, 0.3mM Na_2_HPO_4_, 6mM Glucose, HEPES, 0.5mM MgCl_2_.6H_2_O, 0.4mM MgSO_4_.7H_2_O, 0.5mM EDTA; without CaCl_2_) by Masterflex digital peristaltic pump (Cole-Parmer, Vernon Hills, Illinois, United States of America). After cutting the inferior vena cava 25ml of Collagenase (Roche, Merck, Burlington, Massachusetts, United States of America) solution (1mg/ml) in HBSS (containing 1mM CaCl_2_) was allowed to pass through the liver at a constant flow rate of 3 ml/min. The pieces of digested liver tissue were then minced in HBSS (containing 1mM CaCl_2_). The resulting suspension was then passed through a 100 μm cell strainer (SPL Life Sciences, Pochon, Kyonggi-do, South Korea) and the filtrate was centrifuged at 50 g for 2 min at 4°C. The cellular pellet was carefully resuspended in William’s Medium E (1X) [Gibco, Thermo Fisher Scientific, Waltham, Massachusetts, United States of America] supplemented with 1% antibiotic antimycotic solution (HiMedia, Mumbai, India) for plating. The adhered hepatocytes were maintained in Hepatocytes Basal Medium with Ultraglutamine1 (Lonza, Basel, Switzerland) in Collagen (Thermo Fisher Scientific, Waltham, Massachusetts, United States of America) pre-coated cell culture plates.

### Cell Culture

HepG2 (Human hepatoma cell line, ATCC, Manassas, Virginia, United States of America) and HEK293A (Human embryonic kidney cell line, Thermo Fisher Scientific, Waltham, Massachusetts, United States of America) were cultured in MEM (Eagle’s minimal essential medium) and DMEM (Dulbecco’s Modified Eagle Medium) [HiMedia, Mumbai, India] respectively, supplemented with 10% Fetal Bovine Serum (Gibco, Thermo Fisher Scientific, Waltham, Massachusetts, United States of America) and 1% antibiotic antimycotic solution containing Penicillin, Streptomycin and Amphotericin [HiMedia, Mumbai, India]. HEK293A cells require an additional 1% Non-Essential Amino Acids. HA-Ub plasmid (Addgene, Watertown, Massachusetts, United States of America), mycDDK-JOSD1 plasmid (# RC201968, Origene, Rockville, Maryland, United States of America), and flag-SOCS1 plasmid (#OHu17289, GenScript, Piscataway, New Jersey, United States of America) were transfected using Lipofectamine 2000 (Thermo Fisher Scientific, Waltham, Massachusetts, United States of America) according to the manufacturer’s protocol.

### Cell harvesting, Protein estimation and preparation

Washed cells were homogenized in lysis buffer containing 50 mM Tris-HCl (pH 7.4), 100 mM NaCl, 1 mM EDTA, 1 mM EGTA, 1% Triton X-100 and 0.5% protease inhibitor cocktail (Merck Millipore, Burlington, Massachusetts, United States of America) and the resultant slurry was centrifuged at 20,000g for 20 min at 4°C. Protein concentration was estimated using Bradford assay (BioRad, Hercules, California, United States of America). 30-60 μg of protein samples were prepared using 1X Laemmli’s buffer (5X stock containing 250 mM Tris-HCl (pH 6.8), 10% SDS, 50% glycerol, 0.1% Bromophenol Blue and 10% β-Mercaptoethanol), heated at 95°C for 10 min, cooled and centrifuged at 12000g for 2 min prior to loading into polyacrylamide gel.

### Western Blot

Proteins were resolved in 10% SDS PAGE and transferred to PVDF membrane (Millipore, Burlington, Massachusetts, United States of America) having a pore size of 0.45 μm. Membranes were washed in 1X TBST (pH 7.6) comprising of Tris Base, Sodium Chloride and 1% Tween 20 (Sigma Aldrich, St. Louis, Missouri, United States), and were blocked with 5% skimmed milk powder. The required primary antibody (JOSD1 [Abcam, Cambridge, United Kingdom], SOCS1 [Abclonal, Woburn, Massachusetts, United States of America], Cleaved Caspase 3 [Cell Signaling Technology, Danvers, Massachusetts, United States of America], PARP [Cell Signaling Technology, Danvers, Massachusetts, United States of America], myc tag [Cell Signaling Technology, Danvers, Massachusetts, United States of America], HA tag [Cell Signaling Technology, Danvers, Massachusetts, United States of America], flag tag [Cell Signaling Technology, Danvers, Massachusetts, United States of America], MCL [Abclonal, Woburn, Massachusetts, United States of America], TRX1 [Abclonal, Woburn, Massachusetts, United States of America] and Actin [Cell Signaling Technology, Danvers, Massachusetts, United States of America]) prepared at 1/1000th dilution with 1X TBST containing 1% Bovine Serum Albumin and 0.04% Sodium Azide was added to the membrane and incubated overnight at 4°C. The next day, the membrane was again washed multiple times with 1X TBST to remove any unbound primary antibody. The membrane was then incubated with anti-rabbit secondary antibody (Thermo Fisher Scientific, Waltham, Massachusetts, United States of America) for 1 h at room temperature and washed again for multiple times with 1X TBST. The membrane was then developed using Clarity™ ECL Western Blotting Substrate (BioRad, Hercules, California, United States of America) and viewed in ChemiDoc MP (BioRad, Hercules, California, United States of America).

### Site-directed mutagenesis

JOSD1 C36A mutant was generated using QuickChange II Site-Directed Mutagenesis Kit (Agilent, Santa Clara, California, United States) following the manufacturer’s protocol. The following primers were designed for the process using the primer design guidelines of the kit: Forward Primer: 5’ cagcgcagggagcttgctgccctccacgccctc 3’; Reverse Primer: 5’ gagggcgtggagggcagcaagctccctgcgctg 3’. PCR amplicons were digested with DpnI enzyme, transformed in DH5α competent cells, plasmids were isolated from selected colonies, and sequenced for confirming the mutation.

### Generation of JOSD1 and JOSD1C36A mutant cell lines

HepG2 cells were transfected with mycDDK-JOSD1 and mycDDK JOSD1 C36A mutant plasmids using Lipofectamine 2000. After 48 h, cells were subjected to 100 μg/μl of Gentamycin (InvivoGen, San Diego, California, United States of America) selection and pooled clones stably expressing the proteins of interest were propagated.

### siRNA

siRNA transfection was carried out using Lipofectamine^TM^ RNAiMax Transfection Reagent (Thermo Fisher Scientific, Waltham, Massachusetts, United States of America). 72 h post transfection cells were processed for western blot analysis. Human JOSD1 siRNA SMARTPOOL (Cat No. L-017674-00-0005) and Human SOCS1 siRNA SMARTPOOL (Cat No. L-011511-00-0005) were purchased from Dharmacon, Horizon Discovery, Waterbeach, United Kingdom.

### DUB Screening

A loss of function screening using Human ON-TARGETplus siRNA Library - Deubiquitinating Enzymes (Horizon Discovery, Waterbeach, United Kingdom; Cat No. G-104705) in the HepG2 cell line was designed. HepG2 cells were seeded in 96 well luminescence and fluorescence compatible plates and transfected with the respective siRNAs for 72 h. 16 h before cell harvesting, proteasomal inhibition was imposed upon the cells with 10 μM of MG132 (Sigma Aldrich, St. Louis, Missouri, United States of America). ApoLive-Glo Multiplex Assay Kit (Promega, Madison, Wisconsin, United States of America) as per manufacturer’s protocol. Cell viability was estimated by fluorescence and caspase 3/7 activity was estimated by luminescence. Two-way normalization was done with both the viability measurements (% live cells) and Caspase 3/7 activity measurements. For both assays, scrambled siRNA transfected cells treated with MG132 was used as control group. To avoid accumulating type 1 errors for the experimental replicates we used normalization by summing all data points in a replicate. Values were eliminated by checking for outliers. Measures accounted for both viability and apoptosis was plotted as ring cluster plot by using the *circlize* package in R.

### Live Dead Assay

LIVE/DEAD®Viability/Cytotoxicity Kit for mammalian cells (Thermo Fisher Scientific, Waltham, Massachusetts, United States of America) was used for staining control as well as JOSD1 overexpressing stable cell line after proteasomal inhibition for 16 h. Leica TCS SP8 STED was used for imaging the cells.

### Caspase 3/7 substrate activity assay

Cells were seeded in confocal dishes and treated with MG132 for 16 h. The caspase 3/7 substrate activity assay (Thermo Fisher Scientific, Waltham, Massachusetts, United States of America) reagents were added and incubated as per manufacturer’s protocol. Cells were then visualised using Leica TCS SP8 STED.

### Cell Rox Assay

Cells were subjected to MG132 treatment for 16 h and generation of cellular reactive oxygen species (ROS) was measured using CellROX^TM^ Deep Red Reagent (Thermo Fisher Scientific, Waltham, Massachusetts, United States of America) following manufacturer’s protocol. ROS production was imaged using Leica TCS SP8 STED microscope.

### Cycloheximide Chase assay

HepG2 cells or primary mouse hepatocytes were treated with either 50 μg/ml of cycloheximide (Calbiochem, Merck Millipore, Burlington, Massachusetts, United States of America) alone or in combination with 5 μM of MG132 for required time periods. Cells were then harvested using protein lysis buffers at different time points and analysed using western blot.

### Immunoprecipitation

500 μg of crude cell lysate was precleared with 10 μl of PureProteome Protein A Magnetic Beads (Merck Millipore, Burlington, Massachusetts, United States of America) for 30 min. The supernatant containing the pre-cleared crude cell lysate was collected by attaching the tubes to a Magna Rack (Merck Millipore, Burlington, Massachusetts, United States of America). This pre-cleared crude cell lysate was then used for an overnight reaction at 4°C for immune complex formation with fresh 10 μl of PureProteome Protein A Magnetic Beads and required amount of primary antibody [anti-myc Tag, anti-flag tag (Cell Signalling Technology, Danvers, Massachusetts, United States of America)]. The next day the immune complexes were retrieved by washing magnetic beads with PBS and 0.1% Triton-X100 thrice to remove non-specific molecules. The eluted immune complex was collected by centrifugation and preceded for western blot.

### Immunofluorescence

Cells were fixed with 4% paraformaldehyde for 15 min, washed with 1X PBS and blocked with serum of the species in which the secondary antibody was raised and Triton-X100 for 1 h at room temperature. Cells were then incubated with anti-myc antibody (1:200 dilution) overnight at 4°C followed by incubation with Goat Anti-Rabbit Alexa Fluor 647 conjugated secondary antibody at a ratio of 1:250 for 2 h. The cells were counterstained with 25 µg/ml HOECKST342 and proceeded for visualisation with Leica TCS SP8 STED microscope.

### Image and Statistical analysis

Confocal images were analyzed in Image J by counting the relevant dots for live or dead cells or caspase positive cells across control and experimental groups. For each experiment, n=4 fields/group were taken into consideration for further calculation and analysis. Densitometry was performed by measuring the protein band intensities obtained in western blots using Image J. Obtained values were normalized with actin. All statistical analysis and graphs were done in GraphPad Prism (8.0.1) using Z statistics with p value cut off <0.05.

## RESULTS

### JOSD1 identified as a crucial regulator of cell death under proteotoxic stress

Perturbation in protein turnover and proteastasis via the ubiquitin proteasome pathway cause significant hepatocellular damage. Here we sought to identify the DUBs responsible for this phenomenon. For this, we used a 96 DUB siRNA library and transfected HepG2 cells with the respective siRNAs followed by proteasome inhibitor MG132 treatment for 16 h. Cell death via caspase 3/7 activation and cell viability were assessed as two antipodal functional readouts (r= −0.53, p=0.0001; Supplementary Fig 1A). While majority of the DUBs did not have noticeable impact on cellular death or viability (Figure 1A), we find a cluster of genes knockdown of which resulted in high viability and low apoptosis indicating the causal candidates (*MYSM1*, *USP20*, *USP4*, *USP47*, *USP25*, and UBTD1) facilitating hepatic proteasomal stress (Figure 1B). In contrast, we found that the depletion of *JOSD1* resulted in a significant increase in apoptosis with concomitant loss of viability suggesting its crucial importance in protecting proteotoxic hepatocellular injury (Figure 1A, B). To validate JOSD1 as a pivotal player in modulating cellular apoptosis under proteotoxic stress, we knocked down JOSD1 using siRNA in HepG2 cells and proteotoxicity was induced with MG132 treatment. Downregulation of JOSD1 did not alter cellular ubiquitylated protein profile (Supplementary Fig 1B) but caused an increase in the expression of cleaved caspase 3 (CC3) and cleaved PARP (Fig 1C) with significantly elevated number of dead cells (Fig 1D, top panel). However, there was no change in the level of reactive oxygen species in JOSD1 depleted cells compared to control under proteasomal inhibition indicating that JOSD1 operates in a ROS independent pathway (Fig 1D, bottom panel). A similar observation was seen in primary mouse hepatocytes where, adenovirus mediated knockdown of JOSD1 caused a robust increase in cell death as evidenced by increased expression of CC3 and cleaved PARP when subjected to proteasomal inhibition (Fig 1E).

**Figure 1.**
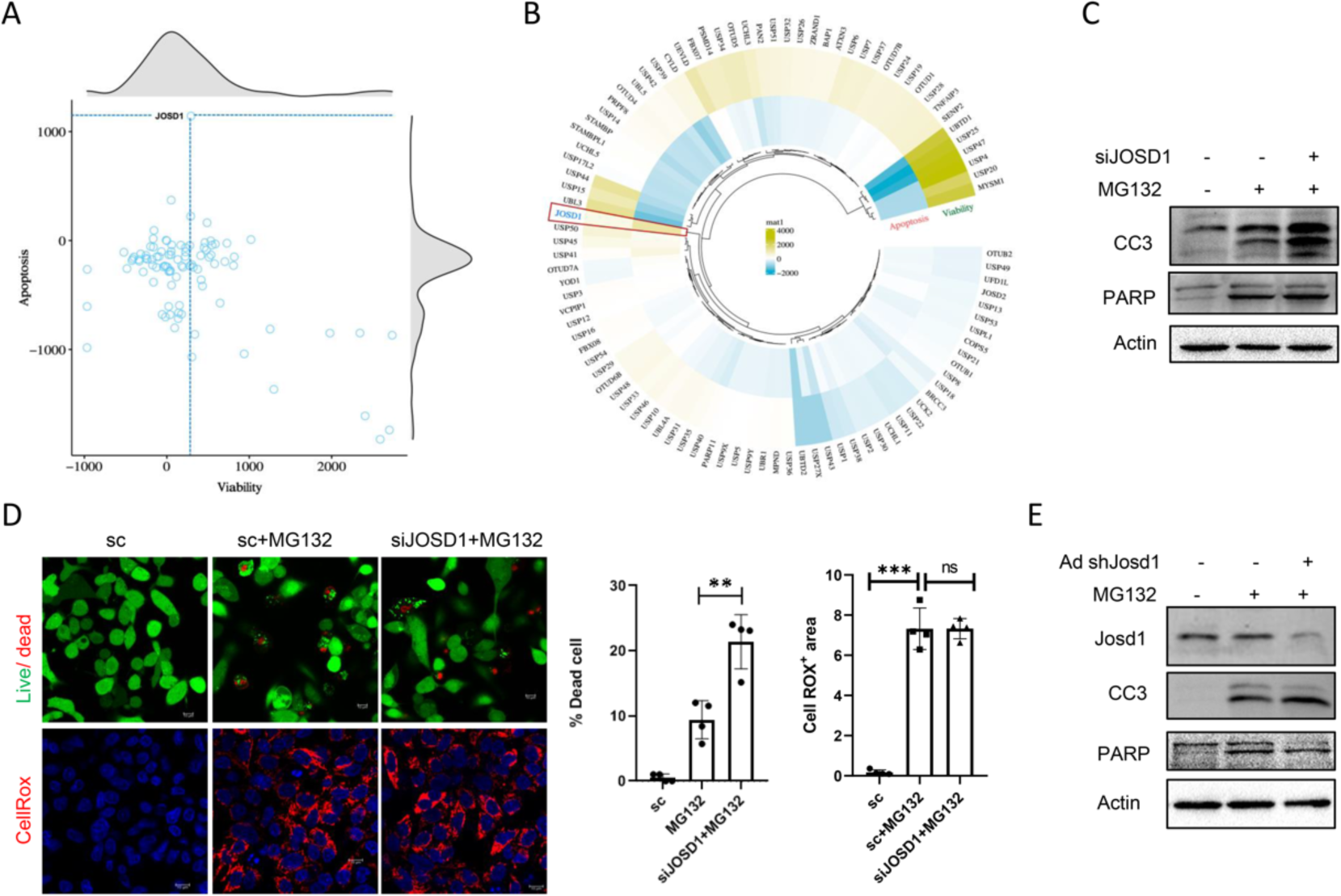
siRNA screening of DUBs and validation of JOSD1. A. 2D scatterplot for the siRNA screening representing relative luminescence (RLU) as a function of apoptosis in the Y-axis, whereas the X-axis indicates relative fluorescence intensity (RFU) as a measure of cell viability. The distribution density for both measures was plotted against the X and Y-axis border. Each point represents a normalized relative intensity from 3 independent experimental replicates. The dotted line (Blue) indicates the respective RFU and RLU values from the JOSD1 knockdown. B. Circular heatmap representing differential clustering in Euclidean distance matrix using the quantitative measures of viability and apoptosis from the siRNA screening. Euclidean clustering groups RFU and RLU values into clusters such that the losses of function for individual genes resulting in response are clustered together. C. JOSD1 knockdown by siRNA transfection in HepG2 cells treated with 5 μM MG132 for 16 h and analyzed for cell death markers. D. (Top) Live Dead Assay in JOSD1 knocked down cells under proteotoxic stress induced with 5μM MG132 for 16 h. Green indicates live cell whereas red dots indicate dead cells. (Bottom) CellRox assay performed in control and JOSD1 depleted cells in proteotoxic condition. Red indicates level of generated cell ROS. (Right) Statistical analysis of percentage of dead cells and CellRox^+^ area. n = 4 fields/group. Values are presented as mean ± SEM. **P < 0.01, ***P < 0.001,. ns - not significant. Scale bar – 10 µM. E. JOSD1 knocked down in primary mouse hepatocytes by adenovirus mediated delivery and checked for apoptosis markers after induction of proteotoxicity by 5 μM MG132 for 16 h.

### JOSD1 confers protection against cell death in proteotoxic conditions

Since absence of JOSD1 caused significant enhancement of apoptosis, we proceeded to address how JOSD1 confers protection to proteotoxic stress. To this end, we made a HepG2 cell line stably expressing myc epitope-tagged JOSD1 for the next set of experiments. In contrast to the depletion of JOSD1, we observed that expression of JOSD1 caused a noticeable reduction in cellular apoptosis as evidenced by decreased expression of CC3 and cleaved PARP (Fig 2A) and in the number of dead cells (Fig 2B). Consistently, adenovirus mediated overexpression of JOSD1 in primary mouse hepatocytes caused a marked decline in cell death under proteotoxicty (Fig 2C). Therefore, these observations serve to affirm that JOSD1 indeed rescues hepatocytes from apoptosis under proteotoxic conditions. Interestingly, we observed an accumulation of JOSD1 proteins following MG132 treatment (Fig 2A), and such accumulation was found to be dependent both with treatment duration (Fig 2D) as well as with the dose of MG132 (Fig 2E). MG132-mediated JOSD1 expression was however not driven by enhanced transcription (Data not shown), but was due to an increase in the half-life of JOSD1 in presence of MG132 as shown by cycloheximide chase assay (Fig 2F). As ubiquitinated JOSD1 could localise in the plasma membrane to influence membrane dynamics, cell motility, and pinocytosis^19^, we sought to investigate if similar translocation and localisation of JOSD1 could be seen under proteasomal inhibition. Immunofluorescence with anti-myc tag antibody showed that JOSD1 was unequivocally accumulated around the cell membrane upon proteasomal inhibition (Fig 2G). Thus, when hepatocytes are exposed to proteotoxic stress, JOSD1 protein is stabilized, accumulated around the plasma membrane and exercises its anti-apoptotic function.

**Figure 2.**
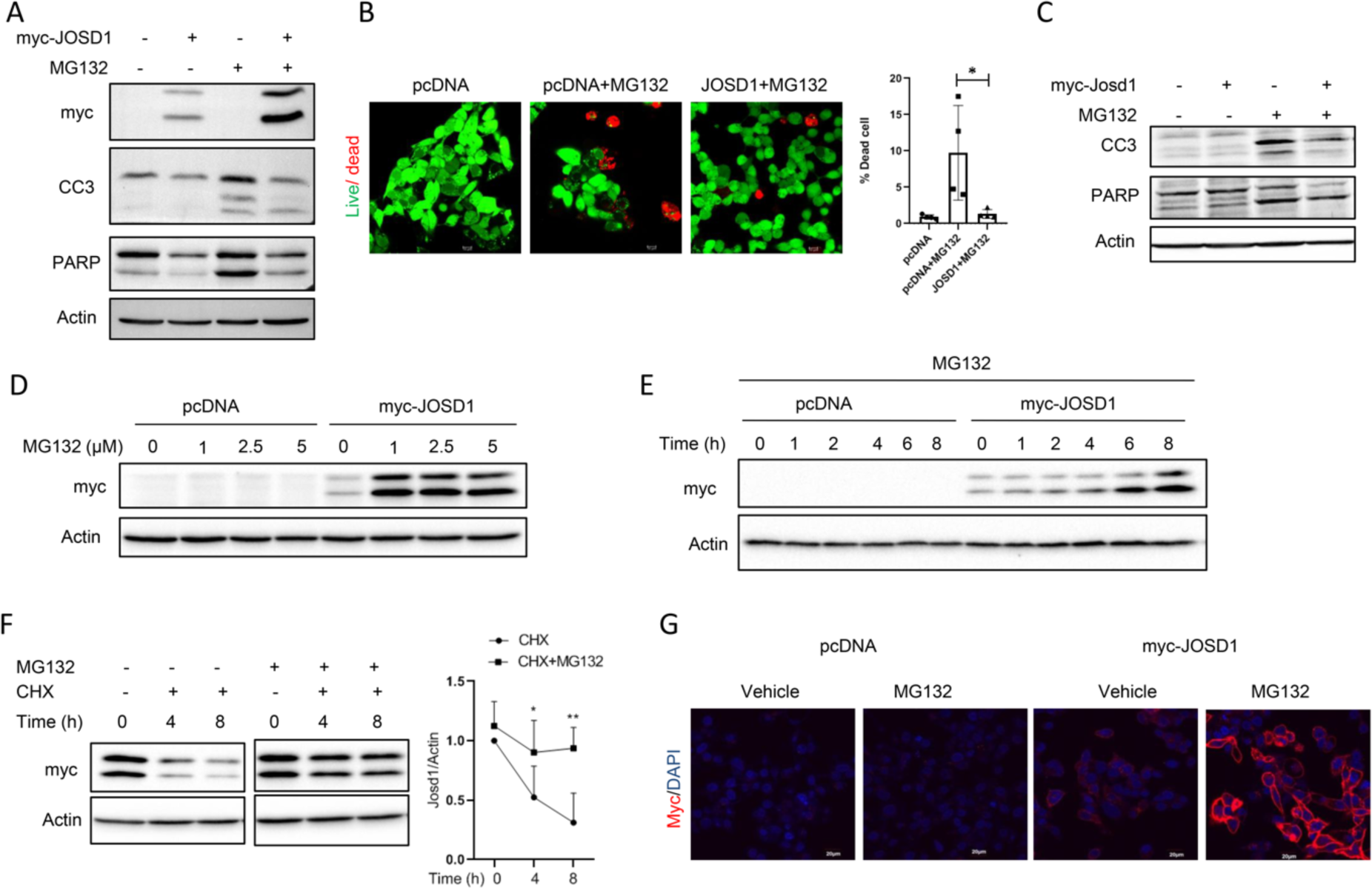
Impact of proteasomal inhibition in HepG2 cells stably expressing JOSD1. A. Status of cellular apoptosis in JOSD1 overexpressing cell line under proteotoxic stress induced by 5 μM MG132 for 16 h. B. Live Dead Assay in JOSD1 overexpression cell line under proteotoxic condition. Green represents live cells and red represents dead cells. (Right) Statistical analysis of percentage of dead cells. n = 4 fields/group. Values are presented as mean ± SEM. *P < 0.05, Scale bar – 10µM. C. JOSD1 overexpressed in primary mouse hepatocytes with the aid of adenovirus mediated delivery and checked for apoptosis markers under proteotoxic stress. D. Accumulation of JOSD1 with increasing dose of MG132 for 16 h in JOSD1 overexpression cell line. E. Accumulation of JOSD1 over time with in JOSD1 overexpressing cell line. F. Cycloheximide chase assay performed under proteotoxic stress. G. Immunofluorescence in JOSD1 overexpressing cell line with anti-myc tag antibody showing JOSD1 cellular localization under proteotoxic stress induced with 5 μM MG132 for 16 h.

### Enzymatically inactive JOSD1 C36A mutant fails to rescue proteotoxicity

JOSD1 has a cysteine residue at 36^th^ position in its active site^20^ and to render the enzyme catalytically inactive, the cysteine was mutated to alanine via site directed mutagenesis. We next made a JOSD1 C36A mutant overexpressing HepG2 cell line. Upon proteasomal inhibition, catalytically inactive enzyme also accumulated over time (Fig 3A), but in contrast to the wild type it was diffusely localised in the cytoplasm (Fig 3B) and failed to mitigate cellular apoptosis (Fig 3C and Supplementary Fig 2A, B). To further establish the requirement of enzymatic activity in protecting proteotoxicity, we performed a caspase 3/7 substrate assay, readout of cell death. Expectedly the caspase 3/7 activity was significantly diminished in JOSD1 overexpressing cells, the activity was restored to that of control cells in JOSD1 C36A mutant cell line under conditions of proteotoxicity (Fig 3D). Since JOSD1 is active when it is monoubiquitinated^19^, we next checked for the difference in the ubiquitination status of wild type JOSD1 and JOSD1 C36A under proteotoxic stress. We found a remarkable difference in the ubiquitination status of the two proteins wherein wild type JOSD1 was predominantly monoubiquitinated and JOSD1 C36A was polyubiquitinated under proteotoxicty (Fig 3E). This observable alteration in the ubiquitination status provides a plausible explanation to the mutant enzyme’s deterred ability to actively exercise protection from cellular apoptosis compared to the wild type protein.

**Figure 3.**
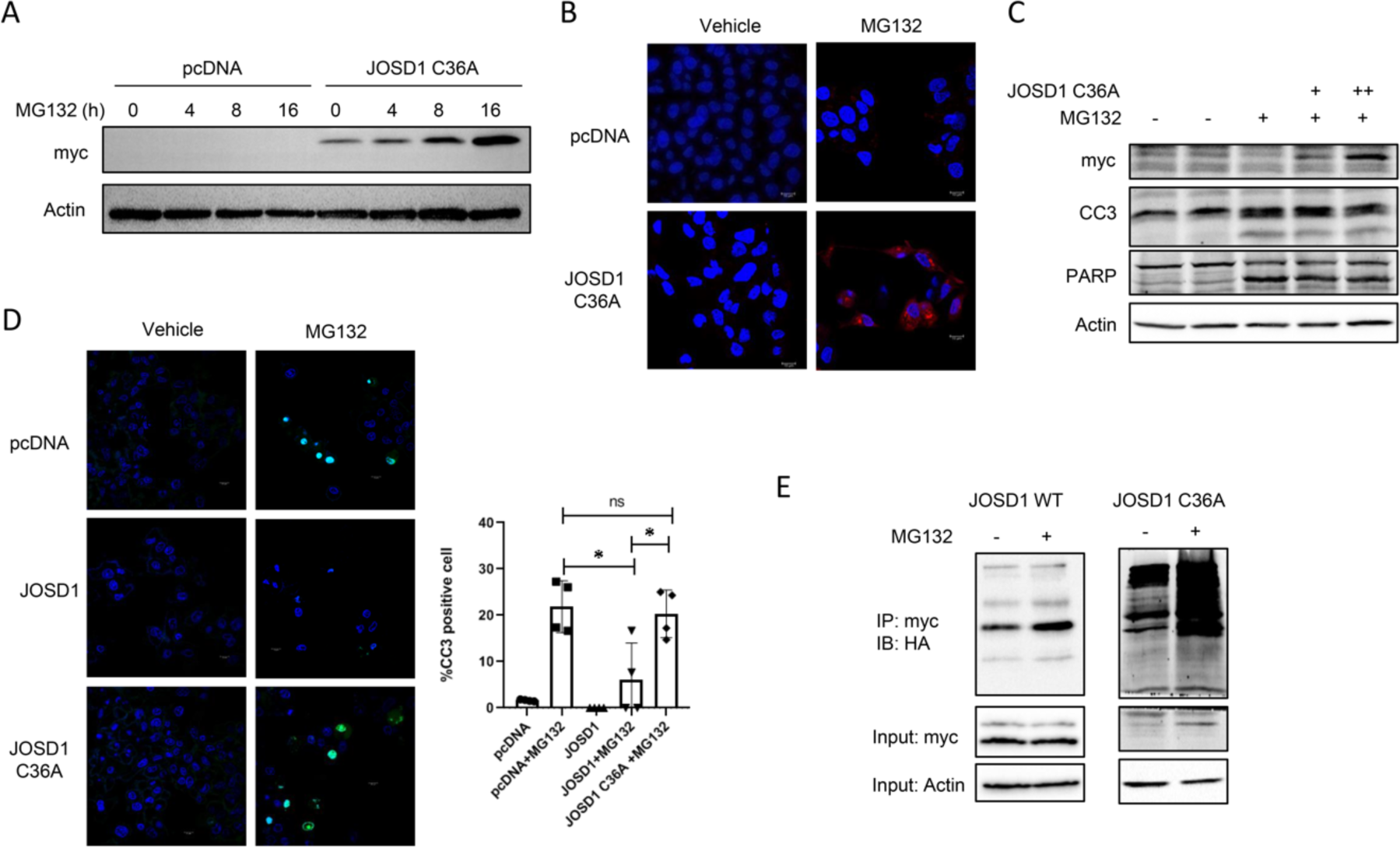
Hepatoprotective effects of JOSD1 C36A mutant during proteotoxicity. A. Accumulation of JOSD1 mutant protein over time with 5 μM MG132 in JOSD1 C36A mutant overexpressing cell line. B. Cellular localization of JOSD1 C36A mutant revealed by immunofluorescence with anti-myc tag antibody in JOSD1 C36A mutant overexpressing cell line under proteotoxic stress. C. Status of hepatic apoptosis in JOSD1 C36A mutant overexpressing cell line under conditions of proteotoxicity. D. (Left) Live caspase activity shown with caspase substrate activity assay in control, JOSD1 WT and JOSD1 mutant overexpressing cell lines during proteotoxicity induced with 5 μM MG132 for 16 h. (Right) Statistical analysis of percentage of cleaved caspase 3 (CC3) positive cells. n = 4 fields/group. Values are presented as mean ± SEM. *P < 0.05. ns - not significant. Scale bar – 10µM. E. HepG2 cells stably expressing myc-tagged wild type JOSD1 or myc-tagged C36A JOSD1 mutant were transfected with HA-tagged ubiquitin. Following Mg132 treatement for 4 h, JOSD1 was immunoprecipitated with anti myc-tag antibody and probed with anti HA-tag antibody.

### JOSD1 interacts with and deubiquitinates SOCS1 to mitigate apoptosis under proteotoxic stress

To decipher how JOSD1 imparts a protective response in proteotoxicity, we sought for its binding partners such as SOCS1 MCL1^20, 21^. Primary hepatocytes treated with MG132 for 16 h revealed an accumulation of JOSD1 and SOCS1 while TRX and MCL1 did not show such increase (Fig 4A) with a noticeable accumulation of SOCS1 in MG132 treated cells (Fig 4B). Since JOSD1 interacts with and deubiquitinates SOCS1 to negatively regulate type I interferon antiviral activity^21^, we sought to identify similar interactions in hepatocytes. Adenovirus mediated ectopic expression of JOSD1 showed a modest increase of SOCS1 levels in primary hepatocytes (Fig 4C). Co-immunoprecipitation assay further show that endogenous SOCS1 interacts with ectopically expressed JOSD1 both in HepG2 and primary mouse hepatocytes (Fig 4D). Interestingly, such interaction is pronounced under proteotoxic condition when expressions of both JOSD1 and SOCS1 are enhanced. Since MG132-mediated enhanced SOCS1 expression was however not transcriptionally regulated (Data not shown), we sought for its protein stability. The half-life of SOCS1 was similarly prolonged in primary mouse hepatocytes in a manner similar to JOSD1 under MG132 treatment (Fig 4E). Moreover, JOSD1 could enhance the half-life of SOCS1 and protect it from cycloheximide mediated decrease with time in HepG2 cells under proteotoxic stress (Fig 4F). Since JOSD1 is a deubiquitinating enzyme, it is expected to deubiquitinate SOCS1 and thus stabilise it. It was indeed seen that JOSD1 could remove ubiquitin chains from SOCS1 and stabilise the protein (Fig 4G). Though JOSD1 C36A mutant is also able to bind SOCS1 (Fig 4H), it does not possess the capability to deubiquitinate SOCS1 under conditions of proteotoxic stress (Fig 4I).

**Figure 4.**
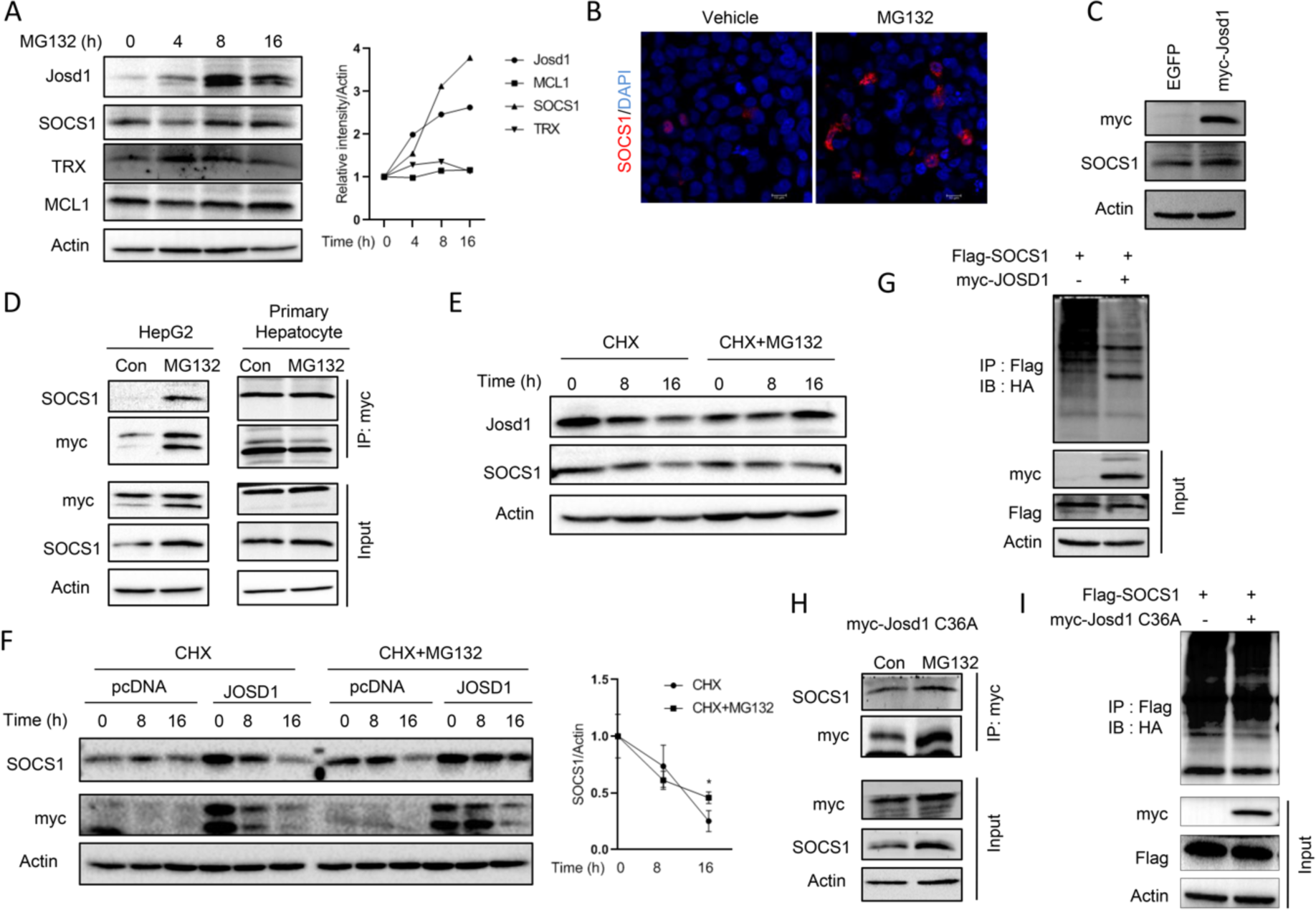
JOSD1 binds, deubiquitinates and stabilizes SOCS1 in hepatocytes. A. (Left) Primary mouse hepatocytes treated with 5 μM MG132 for indicated time periods and checked for JOSD1, SOCS1, TRX and MCL1 protein levels. (Right) Relative band intensities of the corresponding proteins from three experimental replicates. B. Immunofluorescence staining of SOCS1 in HepG2 cells treated with MG132. C. SOCS1 protein expression levels in primary mouse hepatocytes overexpressing JOSD1 by adenovirus mediated method when induced with 5μM MG132 for 16 h. D. Co-Immunoprecipitation of SOCS1 and JOSD1 in JOSD1 overexpressing cells (left) and primary mouse hepatocytes expressing JOSD1 (right) induced with 5μM MG132 for 4 h. E. Cycloheximide (CHX) chase assay of JOSD1 and SOCS1 in primary mouse hepatocytes under proteotoxic stress. F. (Left) Cycloheximide (CHX) chase assay of SOCS1 in JOSD1 overexpressing HepG2 cell line under proteotoxic stress. (Right) Densitometry of band intensities representing half-life of SOCS1 in JOSD1. G. HepG2 cells stably expressing myc-JOSD1 were transfected with Flag-SOCS1 and HA-ubiquitin and immunoprecipitated with anti-Flag antibody following MG132 treatment for 4 h. H. Co-immunoprecipitation of endogenous SOCS1 and JOSD1 in JOSD1 C36A mutant overexpressing cells when induced with 5 μM MG132 for 4 h. I. HepG2 cells stably expressing myc-JOSD1 C36A mutant were transfected with Flag-SOCS1 and HA-ubiquitin and immunoprecipitated with anti-Flag antibody following MG132 treatment for 4 h. Ubiquitination status of SOCS1 shown upon treatment with 5 μM MG132 for 4 h.

### JOSD1 was dependent on SOCS1 to mediate its anti-apoptotic property

Since JOSD1 deubiquitinates and stabilizes SOCS1, we next examined whether SOCS1 alone could impede proteotoxicity in hepatocytes. We find that overexpression of SOCS1 caused a decrease in CC3 expression and number of dead cells (Supplementary Fig 3A-C) while its downregulation conversely caused an increase in CC3 expression (Fig 5A) and number of dead cells (Fig 5B) under proteasomal inhibition in a manner similar to JOSD1. Next we sought to determine whether SOCS1 sufficiency was prerequisite for the JOSD1’s ability to modulate cell death under proteotoxic conditions. To this end, SOCS1 was knocked down in both wild type and mutant JOSD1 overexpressing cells. Absence of SOCS1 attenuated JOSD1’s ability to reverse apoptosis under proteotoxic stress while JOSD1 C36A cells were unaffected (Fig 5C-E). Conversely, overexpression of SOCS1 in JOSD1 depleted cells rescued hepatic cells from undergoing enhanced apoptosis subjected to proteotoxicity (Fig 5F-H). These observations thereby establish SOCS1 as necessary and sufficient downstream effector that aids JOSD1 in shielding hepatocytes from unregulated apoptosis under proteotoxic stress.

**Figure 5.**
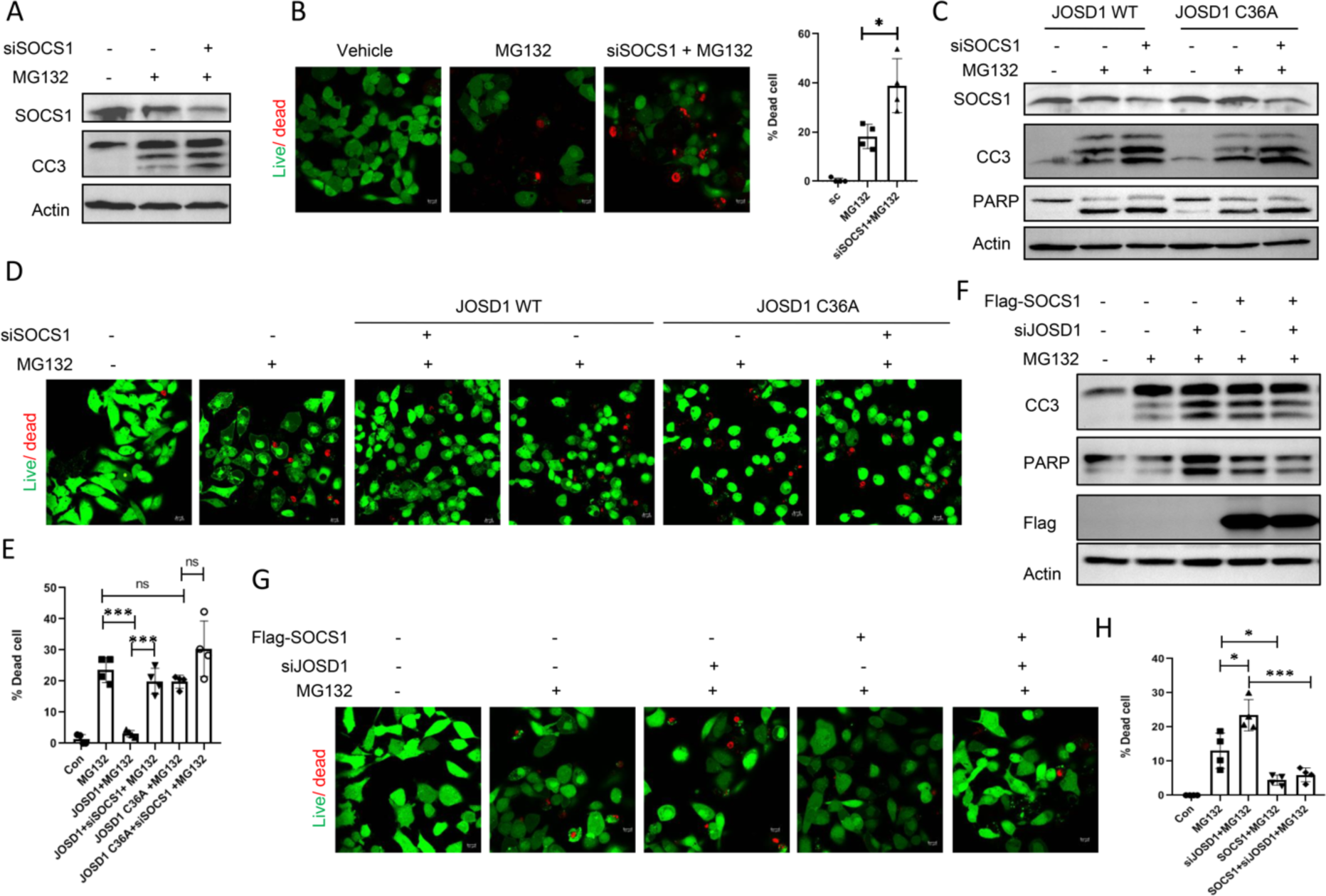
Impact of SOCS1 knockdown and overexpression on the hepatoprotective effects of JOSD1 during proteotoxic stress. A. Expression levels of apoptotic markers in SOCS1 depleted (siSOCS1) HepG2 cells under proteotoxicity induced with 5 μM MG132 for 16 h. B. (left) Live Dead Assay in SOCS1 depleted HepG2 cells under proteotoxic condition. Green represents live cells and red represents dead cells. (Right) Statistical analysis of percentage of dead cells. n = 4 fields/group. Values are presented as mean ± SEM. *P < 0.05, ns - not significant. Scale bar – 10µM. C. Status of expression of apoptotic markers in SOCS1 depleted HepG2 cells stably expressing wild type JOSD1 and JOSD1 C36A mutant under proteotoxicity. D, E. Live Dead Assay in SOCS1 depleted cells stably expressing wild type JOSD1 and JOSD1 C36A mutant under proteotoxic condition. (E) Statistical analysis of percentage of dead cells for Figure 5D. n = 4 fields/group. Values are presented as mean ± SEM. ***P < 0.001. ns - not significant. Scale bar – 10µM. F. HepG2 cells were transfected with Flag-tagged SOCS1 and depleted with JOSD1 (siJOSD1) and levels of apoptosis markers determined by immunoblot. G, H. Live Dead Assay in JOSD1 depleted and SOCS1 expressing HepG2 cells under proteotoxic condition triggered by 5μM MG132 for 16 h. Green represents live cells and red represents dead cells. (H) Statistical analysis of percentage of dead cells for Figure 5G. n = 4 fields/group. Values are presented as mean ± SEM. *P < 0.05, ***P < 0.001. ns - not significant. Scale bar – 10µM.

### JOSD1 ameliorates apoptosis in liver under proteotoxicity

We next examined whether JOSD1 could temper hepatic proteotoxicity in an *in vivo* murine model subjected to acute proteasomal inhibition. To this end, we either knocked down or overexpressed JOSD1 in the livers via adenovirus mediated delivery by tail vein injection. After 7 days, the mice were intraperitoneally injected with 5 mg/kg of bortezomib for 16 h following which they were sacrificed (Fig 6A). Although overexpression of JOSD1 did not alter SOCS1 protein levels (Fig 6B), we find a decline in SOCS1 following knockdown of JOSD1 (Fig 6C). Interestingly, the levels of plasma AST, a liver enzyme which serves as a marker for liver injury, was reciprocally increased upon JOSD1 knockdown and decreased by overexpression of JOSD1 (Fig 6D). However, there were no remarkable changes in liver tissue architecture (H&E), levels of oxidative stress (4-HNE) and ubiquitination status across the three groups (Fig 6E). Notably, knockdown of JOSD1 exacerbated apoptosis as evidenced by increased expression of CC3 both in western blot (Fig 6C) and immunofluorescence assays (Fig 6E). In contrast, ectopic expressions of JOSD1 substantially assuage hepatic injury by lessening cell death compared to mice injected with vehicle controls (Fig 6E). Taken together, JOSD1 serve as a potent hepatoprotective enzyme *in vivo* under proteotoxic stress.

**Figure 6.**
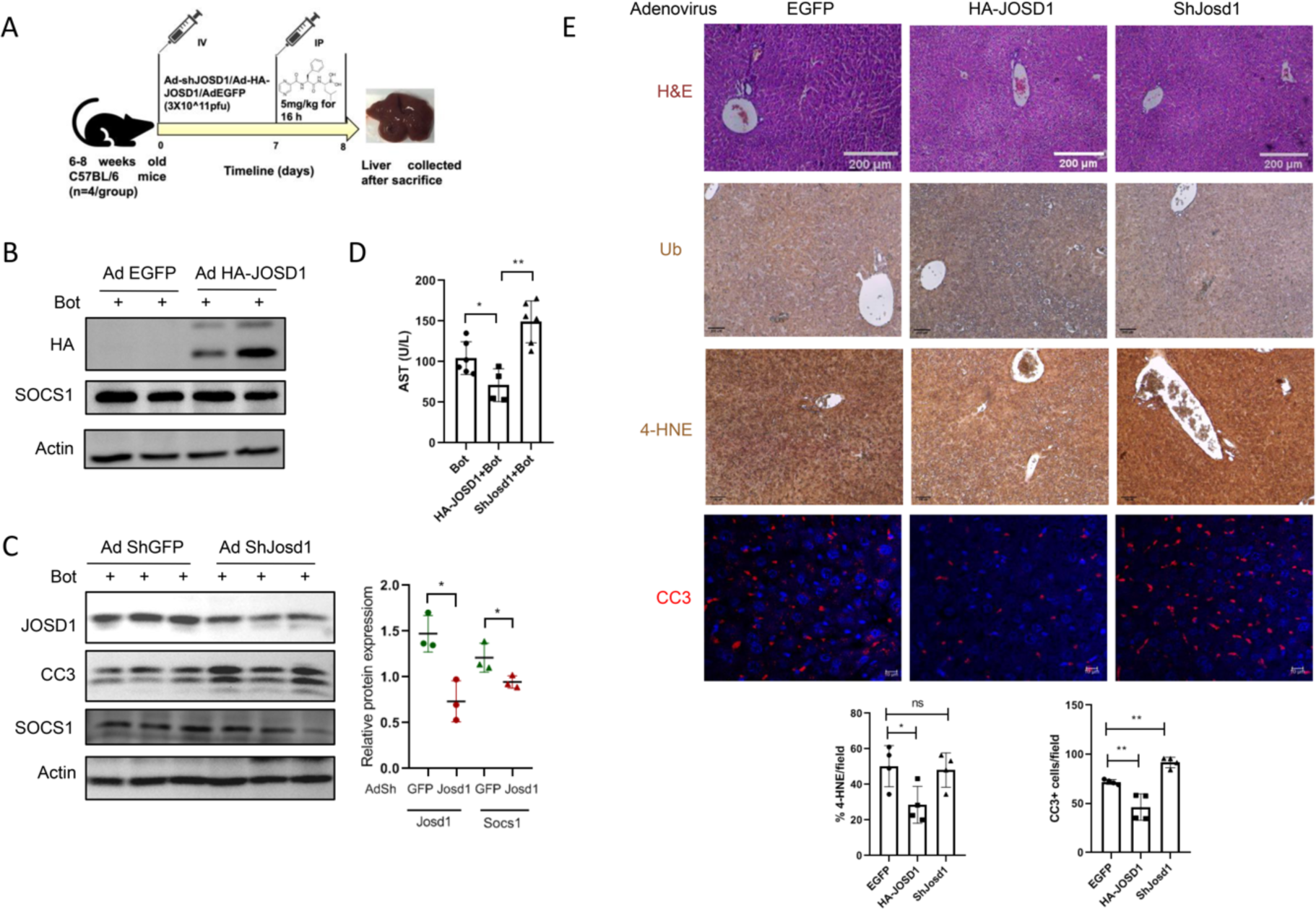
Depletion and ectopic expression of JOSD1 in mouse liver under bortezomib treatment. A. Graphical representation of *in vivo* model and experimental scheme. B. Overexpression of JOSD1 by intravenous injection of 3X10^11^ pfu Ad HA-JOSD1 and status of SOCS1 expression in mouse liver following intraperitoneal administration of 5 mg/kg of bortezomib for 16 h. Control group were injected with 3X10^11^ pfu Ad EGFP. C. (Left) JOSD1 gene was depleted by administering 3X10^11^ pfu Ad shJOSD1 and proteotoxicity was induced by bortezomib. (Right) Statistical analysis showing densitometric analysis of JOSD1 and SOCS1 protein expressions. n = 3 animals/group. Values are presented as mean ± SEM. *P < 0.05, **P < 0.01. ns - not significant. D. Plasma levels of AST in control, JOSD1 knockdown and JOSD1 overexpression murine models of acute proteotoxicity. E. (Top Panels) Liver tissue sections from EGFP, HA-JOSD1 and shJOSD1 carrying adenovirus injected mouse stained with H&E for tissue architecture, immunostained with ubiquitin antibody (Ub), 4-hydroxynonenal (4-HNE) and cleaved caspase 3 (CC3). (Bottom Panels) Statistical analysis of 4-HNE positive area and number of CC3 positive cells across the three groups of animals. n = 4 animals/group. Values are presented as mean ± SEM. *P < 0.05, **P < 0.01. ns - not significant.

## DISCUSSION

The roles and functional implications of DUBs in various cellular events are recently being unravelled. Although accumulating evidence suggested involvement of DUBs in steatotic liver diseases, very few studies, so far, have elucidated the involvement of DUBs in hepatic manifestations of xenobiotic toxicity. Since liver is one of the most proteostatically challenged organs and there are compelling evidences of marked aberrations in protein homeostasis culminating in extensive hepatic injury^2,3^, involvement of UPS and its components in such contexts are unknown. In this study, we explored the roles of DUBs in proteasomal inhibitor-induced models of hepatocellular injury. A first of its kind siRNA screening assay identified JOSD1 as a candidate DUB that reversed hepatic proteotoxicity. We have further identified the downstream molecular event of deubiquitination of SOCS1 through which JOSD1 elicits the protective response. Loss and gain of function studies further established its potential hepatoprotective effects in the *in vivo* setting (Figure 7).

**Figure 7.**
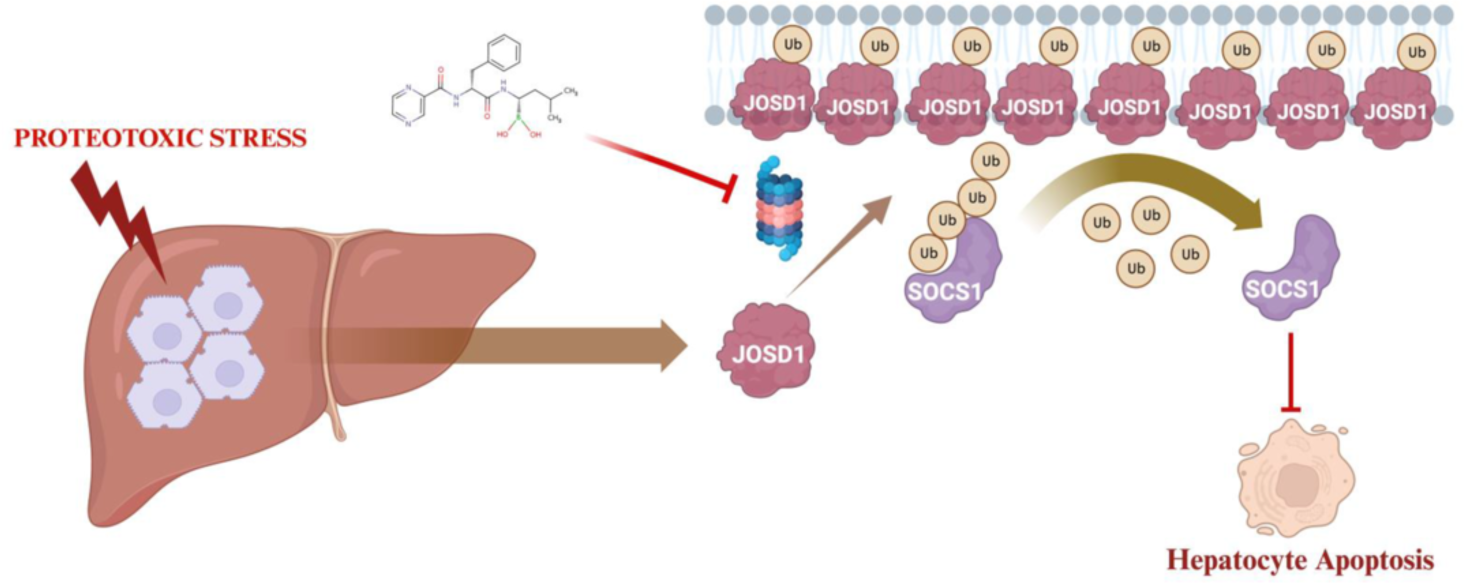
Schematic of the hepatoprotective mechanism of JOSD1.

Under proteotoxic stress, hepatocellular JOSD1 revealed unique molecular features such as monoubiquitination, accumulation and plasma membrane localization which are consistent with the existing literature from different cellular contexts^19,20^. Interestingly, the catalytically inactive JOSD1 mutant was not only ineffective against proteotoxicity, but also did not exhibit such molecular events. However, how exactly monoubiquitinated and membrane bound JOSD1 elicits its protective response needs further investigations. Our study opens a plethora of avenues to consider the question of therapeutic scope and viability of JOSD1 during periods of dysfunctional protein homeostasis. It also needs to be seen how JOSD1 is implicated in chronic liver diseases where there is extensive compromised liver function, hepatic injury and deranged proteostasis.

Towards identifying the downstream JOSD1 effecter responsible for its hepatoprotective effects, study of existing JOSD1 substrates revealed analogous accumulation of SOCS1 under proteotoxic stress. Moreover, SOCS1 also prevents cellular death under proteotoxicity and is sufficient for the protective response of JOSD1. As a negative regulator of type 1 interferon signalling and inflammation, SOCS1 has been shown to modulate cell death in cancer and in acute stress^28,29^. Conversely, overexpression of SOCS1 could augment hepatic steatosis and insulin resistance^30^. SOCS1 at the intersection of metaflammation and infection induced inflammation could thereby balance a tissue specific disease outcome. Beyond the consensus anti-inflammatory function, present study thus describes a new functional role of SOCS1 in the context of hepatic proteotoxicity.

In summary, we have substantially unearthed that JOSD1 bestows hepatocytes with protective armour to shelter cells from apoptosis by increased expression and localisation around plasma membrane ultimately culminating in deubiquitination and stabilising SOCS1. Our data uniquely integrates and assimilates the little antecedent knowledge about the mechanisms of JOSD1 in various cellular events into one consolidated form while also unravelling an intricate involvement of JOSD1 in a previously unknown cellular event, hepatic apoptosis and hepatic injury under conditions of proteotoxic stress.

## Acknowledgements

We thank Sounak Bhattacharya and Rabin Pramanik for assisting with confocal microscopy and animal experiments, respectively.

## Author contributions

SC and AS performed most of the cell based and animal experiments and responsible for acquisition of data. DD and SC conducted siRNA screen and analyzed the data. PC conceptualized and supervised the project and acquired research funds. SC and PC analyzed the data and wrote the paper.

## REFERENCES

1. Zatloukal K, French SW, Stumptner C, Strnad S, Harada M, Toivola DM, et al. From Mallory to Mallory-Denk bodies: what, how and why? Exp Cell Res. 2007;313(10):2033–49.

2. Otoda T, Takamura T, Misu H, Ota T, Murata S, Hayashi H, et al. Proteasome dysfunction mediates obesity-induced endoplasmic reticulum stress and insulin resistance in the liver. Diabetes. 2013;62(3):811–24.

3. Das D, Paul A, Lahiri A, Adak M, Maity SK, Sarkar A, et al. Proteasome dysfunction under compromised redox metabolism dictates liver injury in NASH through ASK1/PPARγ binodal complementary modules. Redox Biol. 2021;45:102043.

4. Kim Y, Kim KY, Lee SH, Chung YY, Yahng SA, Lee SE, et al. A Case of Drug-Induced Hepatitis due to Bortezomib in Multiple Myeloma. Immune Netw. 2012;12(3):126–8.

5. Kouroukis TC, Baldassarre FG, Haynes AE, Imrie K, Reece DE, Cheung MC. Bortezomib in multiple myeloma: systematic review and clinical considerations. Curr Oncol. 2014;21(4):e573–603.

6. Rosiñol L, Montoto S, Cibeira MT, Bladé J. Bortezomib-induced severe hepatitis in multiple myeloma: a case report. Arch Intern Med. 2005;165(4):464–5.

7. Cornelis T, Beckers EA, Driessen AL, van der Sande FM, Koek GH. Bortezomib-associated fatal liver failure in a haemodialysis patient with multiple myeloma. Clin Toxicol (Phila). 2012;50(5):444–5.

8. Koschny R, Ganten TM, Sykora J, Haas TL, Sprick MR, Kolb A, et al. TRAIL/bortezomib cotreatment is potentially hepatotoxic but induces cancer-specific apoptosis within a therapeutic window. Hepatology. 2007;45(3):649–58.

9. Lam YA, Xu W, DeMartino GN, Cohen RE. Editing of ubiquitin conjugates by an isopeptidase in the 26S proteasome. Nature. 1997;385(6618):737–40.

10. Pickart CM, Rose IA. Ubiquitin carboxyl-terminal hydrolase acts on ubiquitin carboxyl-terminal amides. J Biol Chem. 1985;260(13):7903–10.

11. Nijman SM B, Luna-Vargas MPA, Velds A, Brummelkamp TR, Dirac AMG, Sixma TK, et al. A genomic and functional inventory of deubiquitinating enzymes. Cell. 2005;123(5):773–86.

12. Hemelaar J, Galardy PJ, Borodovsky A, Kessler BM, Ploegh HL, Ovaa H. Chemistry-based functional proteomics: mechanism-based activity-profiling tools for ubiquitin and ubiquitin-like specific proteases. J Proteome Res. 2004;3(2):268–76.

13. Amerik AY, Hochstrasser M. Mechanism and function of deubiquitinating enzymes. Biochim Biophys Acta. 2004;1695(1-3):189–207.

14. Ji YX, Huang Z, Yang X, Wang X, Zhao LP, Wang PX, et al. The deubiquitinating enzyme cylindromatosis mitigates nonalcoholic steatohepatitis. Nat Med. 2018;24(2):213–223.

15. Liu D, Zhang P, Zhou J, Liao R, Che Y, Gao MM, et al. TNFAIP3 Interacting Protein 3 Overexpression Suppresses Nonalcoholic Steatohepatitis by Blocking TAK1 Activation. Cell Metab. 2020;31(4):726–740.

16. Zhang P, Wang PX, Zhao LP, Zhang X, Ji YX, Zhang XJ, et al. The deubiquitinating enzyme TNFAIP3 mediates inactivation of hepatic ASK1 and ameliorates nonalcoholic steatohepatitis. Nat Med. 2018;24(1):84–94.

17. An S, Zhao LP, Shen LJ, Wang S, Zhang K, Yu Q, et al. USP18 protects against hepatic steatosis and insulin resistance through its deubiquitinating activity. Hepatology. 2017;66(6):1866–1884.

18. Molusky MM, Li S, Ma D, Yu L, Lin JD. Ubiquitin-specific protease 2 regulates hepatic gluconeogenesis and diurnal glucose metabolism through 11β-hydroxysteroid dehydrogenase 1. Diabetes. 2012;61(5):1025–1035.

19. Seki T, Gong L, Williams AJ, Sakai N, Todi SV, Paulson HL. JOSD1, a membrane-targeted deubiquitinating enzyme, is activated by ubiquitination and regulates membrane dynamics, cell motility, and endocytosis. J Biol Chem. 2013;288(24):17145–55.

20. Wu X, Luo Q, Zhao P, Chang W, Wang Y, Shu T, et al. JOSD1 inhibits mitochondrial apoptotic signalling to drive acquired chemoresistance in gynaecological cancer by stabilizing MCL1. Cell Death Differ. 2020;27(1):55–70.

21. Wang X, Zhang L, Zhang Y, Zhao P, Qian L, Yuan Y, et al. JOSD1 Negatively Regulates Type-I Interferon Antiviral Activity by Deubiquitinating and Stabilizing SOCS1. Viral Immunol. 2017;30(5):342–349.

22. Jing C, Liu D, Lai Q, Li L, Zhou M, Ye B, et al. JOSD1 promotes proliferation and chemoresistance of head and neck squamous cell carcinoma under the epigenetic regulation of BRD4. Cancer Cell Int. 2021;21(1):375.

23. Yang J, Weisberg EL, Liu X, Magin RS, Chan WC, Hu B, et al. Small molecule inhibition of deubiquitinating enzyme JOSD1 as a novel targeted therapy for leukemias with mutant JAK2. Leukemia. 2022;36(1):210–220.

24. Ma X, Qi W, Yang F, Pan H. Deubiquitinase JOSD1 promotes tumor progression via stabilizing Snail in lung adenocarcinoma. Am J Cancer Res. 2022;(5):2323–2336.

25. Huang S, Huang J, Cui M, Wu X, Wang M, Zhu D, et al. Duck Tembusu Virus Inhibits Type I Interferon Production through the JOSD1-SOCS1-IRF7 Negative-Feedback Regulation Pathway. J Virol. 2022;96(18):e0093022.

26. Li M, Gao F, Li X, Gan Y, Han S, Yu X, et al. Stabilization of MCL-1 by E3 ligase TRAF4 confers radioresistance. Cell Death Dis. 2022;13(12):1053.

27. Mao T, Shao M, Qiu Y, Huang J, Zhang Y, Song B, et al. PKA phosphorylation couples hepatic inositol-requiring enzyme 1alpha to glucagon signaling in glucose metabolism. Proc Natl Acad Sci USA. 2011;108(38):15852–7.

28. Zhang L, Xu C, Ma Y, Zhu K, Chen X, Shi Q, et al. SOCS-1 ameliorates smoke inhalation-induced acute lung injury through inhibition of ASK-1 activity and DISC formation. Clin Immunol. 2018;191:94–99.

29. Beaurivage C, Champagne A, Tobelaim WS, Pomerleau V, Menendez A, Saucier C. SOCS1 in cancer: An oncogene and a tumor suppressor. Cytokine. 2016;82:87–94.

30. Ueki K, Kondo T, Tseng YH, Kahn CR. Central role of suppressors of cytokine signaling proteins in hepatic steatosis, insulin resistance, and the metabolic syndrome in the mouse. Proc Natl Acad Sci USA. 2004;101(28):10422–7.

