## Supplementary material for "Deubiquitinase JOSD1 tempers hepatic proteotoxicity": SUPPLEMTENTARY INFORMATION

**Supplementary Information**

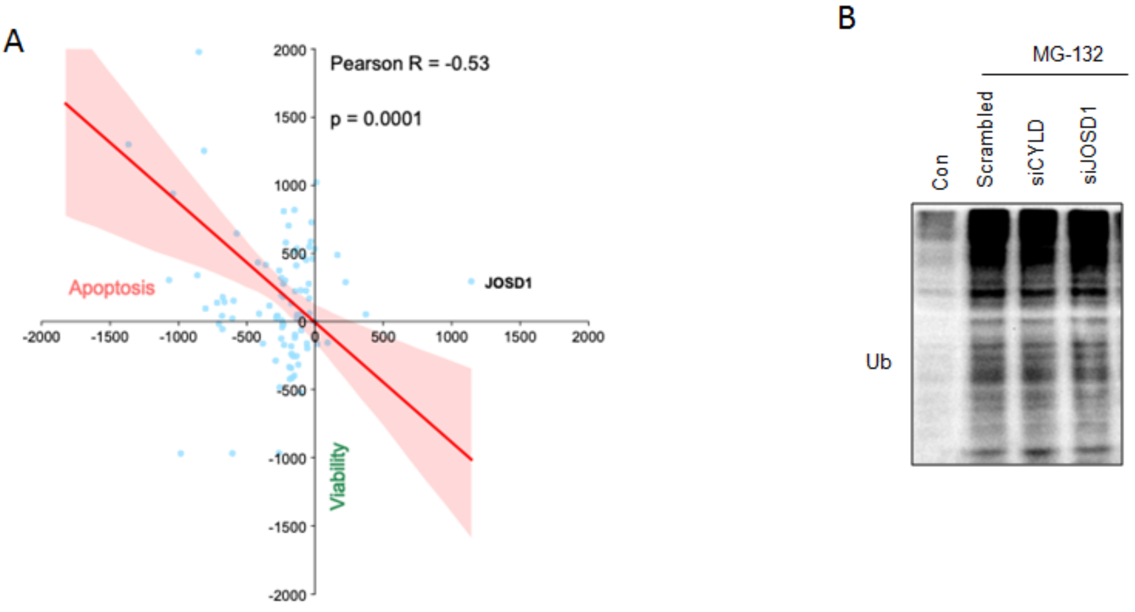


Supplementary Figure1: A. The joint analysis scatter plot represented two variables apoptosis and viability in two dimensions for individual genes in the siRNA screen. Tested for Pearson's correlation and Wilcoxon test for significance testing. Pearson's line is indicated in red and individual points are in blue. B. Total ubiquitinatation in HepG2 cells after knocking down DUBs CYLD and JOSD1 under 10µM MG132 treatment for 16 h.

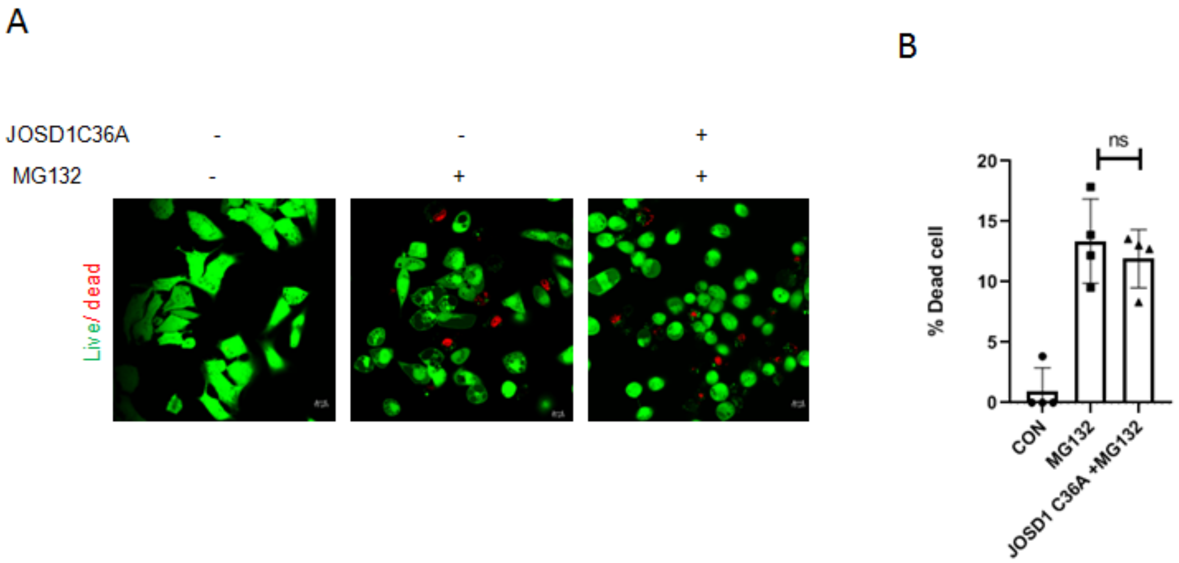


Supplementary Figure 2: A. Live Dead Assay performed in JOSD1C36A mutant cell line after 5µM MG132 treatment for 16 h. B. Graph representing the percentage of dead cells across groups in the live dead assay.

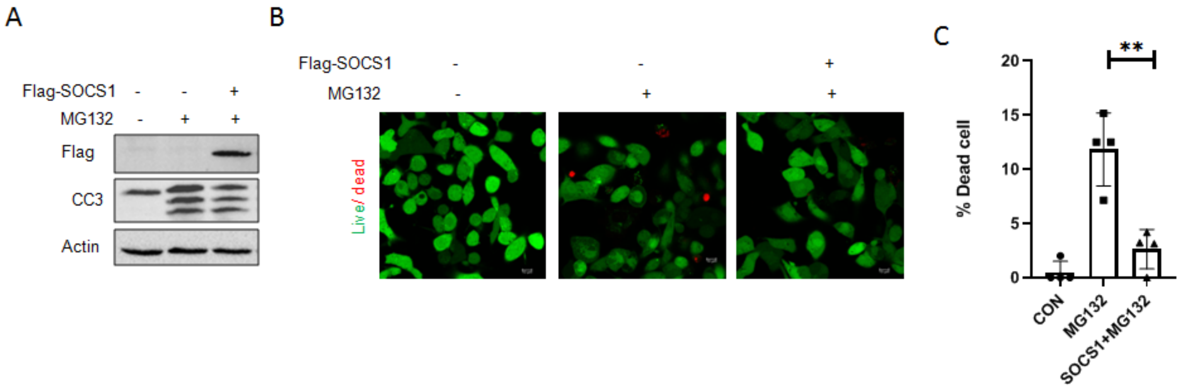


Supplementary Figure 3: A. Western Blot showing cleaved caspase 3 (CC3) levels in HepG2 cells overexpressing SOCS1 under 5μM MG132 treatment for 16 h. B. Live Dead assay in HepG2 cells overexpressing SOCS1 under 5μM MG132 treatment for 16 h. C. Graph representing the percentage of dead cells across groups in the live dead assay.
